# Exact polynomial-time isomorphism testing in directed graphs through comparison of vertex signatures in Krylov subspaces

**DOI:** 10.1101/2022.07.28.501884

**Authors:** Robert John O’Shea

## Abstract

**Motivation:** The complexity of the isomorphism problem in directed graphs has remained unsolved to date. This study examines the properties of Krylov matrices, demonstrating that they may be used to generate vertex “signatures” to allow exact analogy testing in directed graphs.

**Results:** A “vertex signature” is defined by initialising a Krylov matrix with a binary vector indicating the vertex position. This study demonstrates that signatures of analogous vertices are related by a linear-ordering transformation. It is demonstrated that equality of ordered vertex signatures is necessary and sufficient to demonstrate analogy. Thus, analogous vertices may be identified by checking each of the *n* candidates sequentially. This result is extended to analogous vertex sets. Thus, the isomorphic mapping may be constructed iteratively ℴ(*n*^5^) time by building a set of vertex analogies sequentially. The algorithm is applied to a dataset of enzyme structures, with comparison to a common heuristic algorithm.

**Availability and Implementation:** Source code is provided at github.com/robertoshea/graph_isomorphism_directed.

**Supplementary Data:** Supplementary results are attached.

## Background

The graph isomorphism problem is an important issue in complexity science^1^, with various applications in bioinformatics, chemoinformatics, database design and imaging. Polynomial-time isomorphism testing in degenerate undirected graphs remained an open problem since at least 1957^1,2^. Recent results have demonstrated that, vertex analogy may be tested in undirected graphs by checking equality of ordered vertex eigenprojections^3^. Subsequent perturbation of analogous vertex pairs facilitates polynomial time isomorphism testing^3^. However, isomorphism of graph eigenprojections does not hold for directed graphs, which have asymmetrical adjacency matrices and potentially complex eigenbases. This analysis presents an alternative approach for vertex analogy testing in undirected graphs, through comparison of Krylov subspaces. It is demonstrated that Krylov subspaces of analogous vertices are related by a linear permutation, and that equality of the ordered subspaces suffices to demonstrate analogy. Thus, isomorphic mappings may be generated in polynomial time, through the identification of analogies for each vertex in the source graph.

### Related Work

Babai and Luks have presented several heuristic algorithms for undirected graphs with bounded degeneracy ^4–7^. Babai proposed spectral assignment for isomorphism testing in undirected graphs with eigenvalue multiplicity^4^. Babai proposed a spectral assignment algorithm for graphs with bounded eigenvalue multiplicity ^4^. Luks proposed an algorithm for graphs of bounded degree ^5^ which Babai extended to provide a quasipolynomial-time solution for the general case ^6,7^.

Klus and Sahai proposed a heuristic vertex analogy test for undirected graphs by comparing vertex “eigenpolytypes”^8^. Subsequent graph perturbation at vertex pairs estimated to be analogous decreased graph degeneracy heuristically, although “backtracking” was required when non-analogous pairs were perturbed, breaking isomorphism^8^. Vertex analogy may be tested exactly in undirected graphs by comparison of ordered vertex eigenprojections^3^, which apply the same permutation to each column. Subsequent perturbation of analogous vertices allows polynomial-time inference of isomorphic mappings. However, equality of graph eigenprojections is not guaranteed in directed graphs, which may have asymmetric adjacency matrices^3^.

Graph Krylov subspaces, which contain products of vectors and exponentiated adjacency matrices, are closely related to the tensor of eigenprojections^9^, presenting an related avenue for isomorphism research. Ramraj and Prabhakar demonstrated relationships between Krylov subspaces of analogous graphs, generating “proliferation matrices” by initialising Krylov subspaces from column vectors of ones^10^, comparing row-sums of exponentiated adjacency matrices. Although isomorphic graphs are mapped to identical signatures^10^, some non-isomorphic graphs also map to identical signatures, such as those associated with the following adjacency matrices:

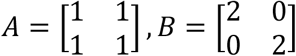

### Preliminaries and notation

Notation follows recent related works^3,8^. Let 𝒮_*n*_ denote the symmetric group of degree *n* and 𝒫_*n*_ denote the set of *n* × *n* dimensional permutation. Let *π* ∈ 𝒮_*n*_ and *P* ∈ 𝒫_*n*_ denote corresponding vector and matrix representations of the same permutation, such that:

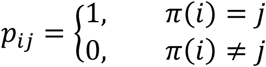

Throughout the article, vector and matrix representation of permutation functions will be used interchangeably, such that *P* ⇔ *π*. Let the ordered set *π*(*C*) = {*π*(*c*_1_), …, *π*(*c*_|*C*|_)} denote the images of the elements of the ordered set *C* ⊆ {1, …, *n*} under permutation *π*.

Let 𝒢_*A*_(𝒱, ℰ_*A*_) and 𝒢_*B*_(𝒱, ℰ_*B*_) be two weighted directed graphs, with asymmetrical adjacency matrices *A, B* ∈ ℝ^*n*×*n*^, where 𝒱 = {*𝓋*_1_, …, *𝓋*_*n*_} is the vertex set and ℰ_*A*_ and ℰ_*B*_ are edge sets. 𝒢_*A*_ and 𝒢_*B*_ are termed the “source” and “target” graphs, respectively. Let 𝒢_*A*_(*𝓋*_*i*_) denote the *i*th vertex in 𝒢_*A*_ and let *A*(*𝓋*_*i*_) denote the corresponding matrix column.

#### Definition.

Graphs 𝒢_*A*_ and 𝒢_*B*_ are termed “isomorphic” if there exists some permutation *π* ∈ 𝒮_*n*_ such that:

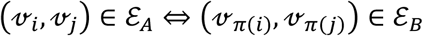

Equivalently, 𝒢_*A*_ and 𝒢_*B*_ are isomorphic if there exists some *P* ∈ 𝒫_*n*_, such that:

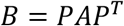

Isomorphism of 𝒢_*A*_ and 𝒢_*B*_ is denoted 𝒢_*A*_ ≅ 𝒢_*B*_.

#### Definition.

A permutation *π* ∈ 𝒮_*n*_ is termed an “isomorphic mapping” from 𝒢_*A*_ to 𝒢_*B*_ if it satisfies:

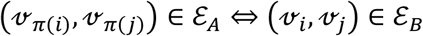

Equivalently, *P* ∈ 𝒫_*n*_ is termed an isomorphic mapping from 𝒢_*A*_ to 𝒢_*B*_ if it satisfies *B* = *PAP*^*T*^. This is denoted *π*: 𝒢_*A*_ → 𝒢_*B*_.

It is emphasised here that the double right arrow “⇒” will be used to denote implication of the right statement by the left., which is distinct from the single left-right arrow “→” used to denote function mappings.

#### Definition.

“Analogy” of the *i*th vertex in 𝒢_*A*_ to the *j*th vertex in 𝒢_*B*_, implies that there exists some 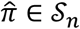 such that 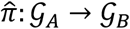 and 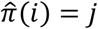. This is denoted 𝒢_*A*_(*𝓋*_*i*_) ↔ 𝒢_*B*_(*𝓋*_*j*_). Likewise, “set analogy” of *𝓋*_*C*_ ⊆ 𝒱 in 𝒢_*A*_ to *𝓋*_*D*_ ⊆ 𝒱 vertex in 𝒢_*B*_, implies that there exists some 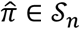 such that 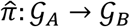 and 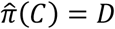. This is denoted 𝒢_*A*_(*𝓋*_*C*_) ↔ 𝒢_*B*_(*𝓋*_*C*_). It is emphasised here that the double left-right arrow “⇔” will be used to denote equivalence of statements, which is distinct from the single left-right arrow “↔” used to denote analogy.

#### Definition.

For some matrix *X* ∈ ℝ^*n*×*n*^, let *Q*_*X*_ ∈ 𝒫_*n*_ be a row-permutation matrix satisfying:

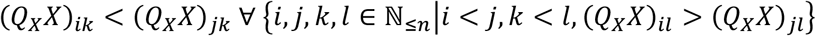

Thus, pre-multiplication by *Q*_*X*_ orders rows in ascension by the first column, then the second, then the third etc, resolving ties in former columns by elements of latter columns. It is noted the solution to *Q*_*X*_ may not be unique if some rows of *X* are identical.

#### Definition.

Let *s*: ℝ^*n*×*m*^ → ℝ^*n*×*m*^ denote the row-ordering function *s*(*X*) = *Q*_*X*_*X*.

#### Definition.

Let *K*(*A, x*) ∈ ℝ^*n*×*n*^ denote the Krylov matrix formed by iterative multiplication of *A* ∈ ℝ^*n*×*n*^ and *x* ∈ ℝ^*n*^, such that:

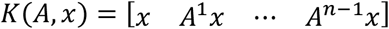

Here *x* is termed the “initialisation vector” of *K*(*A, x*). Let *K*^⊥^(*A, x*) denote an orthogonalisation of the Krylov matrix^11^ by column-wise Arnoldi iteration^12^ such that:

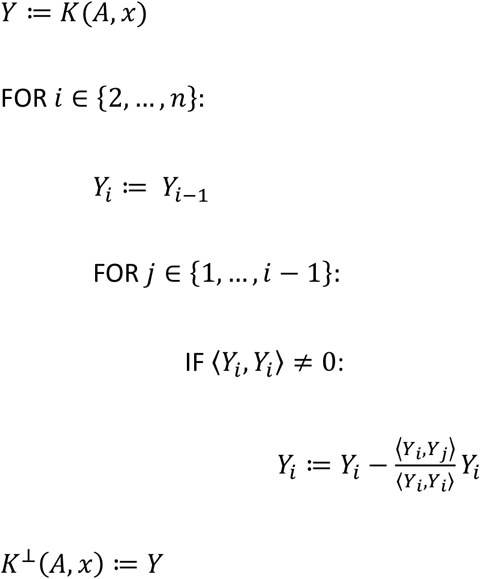

#### Definition.

Let *z*^{*i*}^ ∈ {0,1}^*n*^ such that

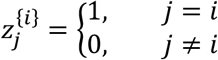

Likewise, let *z*^{*C*}^ ∈ {0,1, …, |*C*|}^*n*^ such that

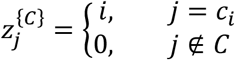

Here *C* is an ordering of the set {1,2, …, |*C*|}, where |*C*| ≤ *n*. It is observed that as 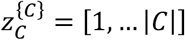. Hence, the elements of 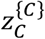 are distinct. It is also noted that all permutations of *C* are distinct.

#### Definition.

The “vertex signature” of the *i*th vertex in 𝒢_*A*_ is the row-ordered, orthogonalized Krylov matrix initialised by the vector *z*^{*i*}^, *K*^⊥^(*A, z*^{*i*}^). Likewise, the row-ordered, orthogonalized Krylov matrix initialised by the vector *z*^{*C*}^, *K*^⊥^(*A, z*^{*C*}^) is termed the “vertex set signature”.

#### Definition.

The “ordered vertex signature” of the *i*th vertex in 𝒢_*A*_ is the row-ordered vertex signature *s* (*K*^⊥^(*A, z*^{*i*}^)). Likewise, *s* (*K*^⊥^(*A, z*^{*C*}^)) is termed the “ordered vertex set signature”.

### Method Overview

This study presents a novel algorithm for isomorphism testing in directed, potentially degenerate graphs, using Krylov matrices. By initialising the Krylov subspace with the vector *z*^{1}^ = [1,0, …, 0], a vertex signature *K*(*A, z*^{1}^) is generated which fully describes the structural identity of 𝒢_*A*_(*𝓋*_1_). This signature is shown to be related to the signatures of analogous vertices by a linear permutation. It is demonstrated that equality of ordered signatures is both necessary and sufficient to demonstrate analogy, providing a polynomial-time test. Following identification of the first analogous vertex pair, a second analogy may be sought by initialising the Krylov subspace with the vector *z*^{1,2}^ = [1,2,0, …, 0]. Fixing the positing of the previously established vertex analogy, only *n* − 1 candidate vertices remain for analogy testing. Thus, analogy testing may proceed until analogies have been identified for each vertex, yielding an isomorphic mapping.

### Theoretical Basis of Algorithm

When searching for an isomorphic mapping between graphs, it is permissible to assume their isomorphism. If an isomorphic mapping is identified, this assumption is validated.

**Lemmas 1a-1d** demonstrate that signatures of analogous vertices are related by a linear permutation, which corresponds to the isomorphic mapping from the source graph to the target. This result provides a necessary condition for vertex analogy testing.

**Lemma 1a** demonstrates that the Krylov subspaces of isomorphic graphs 𝒢_*A*_ and 𝒢_*B*_, where *B* = *PAP*^*T*^ are related by a linear permutation corresponding to their isomorphic mapping, such that:

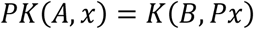

#### Lemma 1a.

If *B* = *PAP*^*T*^ then *PK*(*A, x*) = *K*(*B, Px*)

*Proof*

Substituting *B* for *PAP*^*T*^ we have:

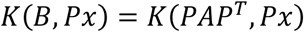

Expanding *K*(*PAP*^*T*^, *Px*) we have:

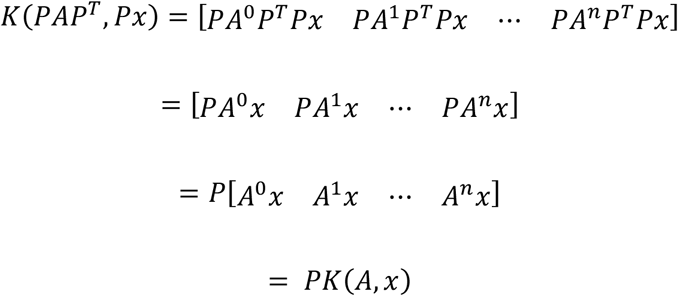

*End of Lemma 1a*

**Lemma 1b** extends the relation described in Lemma 1a to the orthogonalized Krylov subspaces.

#### Lemma 1b.

If *B* = *PAP*^*T*^ then *PK*^⊥^(*A, x*) = *K*^⊥^(*B, Px*)

*Proof*

Let *W* ∈ ℝ^*n*×*n*^ such that:

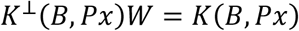

Substituting *B* for *PAP*^*T*^ we have:

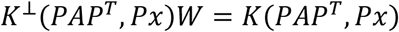

Thus,

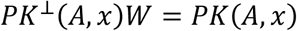

From lemma 1a we have *PK*(*A, x*) = *K*(*B, Px*). Thus, we have:

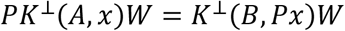

Substituting the definitions of *PK*^⊥^(*A, x*) and *K*^⊥^(*B, Px*), we have:

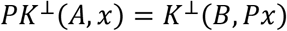

*End of Lemma 1b*

**Lemma 1c** demonstrates that the signatures of analogous vertices are related by the same linear permutation which maps the graphs to one another.

#### Lemma 1c

If *B* = *PAP*^*T*^ then *PK*^⊥^(*A, z*^{*i*}^) = *K*^⊥^(*B, z*^{*π*(*i*)}^).

*Proof*

As *z*^{*π*(*i*)}^ = *Pz*^{*i*}^ we have *K*^⊥^(*B, z*^{*π*(*i*)}^) = *K*^⊥^(*B, Pz*^{*i*}^). By lemma 1b this is true.

*End of lemma 1c*

**Lemma 1d** demonstrates that the signatures of analogous vertex sets are related by same linear permutation which maps the source graph to the target graph.

#### Lemma 1d

If *B* = *PAP*^*T*^ then *PK*^⊥^(*A, z*^{*C*}^) = *K*^⊥^(*B, z*^{*π*(*C*)}^)*Proof*

As *z*^{*π*(*C*)}^ = *Pz*^{*C*}^ we have *K*^⊥^(*B, z*^{*π*(*C*)}^) = *K*^⊥^(*B, Pz*^{*C*}^). By lemma 1b this is true.

*End of lemma 1d*

**Lemma 2** demonstrates that the linear ordering function *s* induces equality between the signatures of analogous vertex sets, so that analogous vertex sets have equal ordered signatures. Therefore, equality of ordered vertex signatures is a necessary condition for vertex set analogy. This result provides a convenient necessary condition for analogy testing.

#### Lemma 2

If *π*: 𝒢_*A*_ → 𝒢_*B*_ and *π*(*i*) = *j* then *s* (*K*^⊥^(*A, z*^{*C*}^)) = *s* (*K*^⊥^(*B, z*^{*π*(*C*)}^)).

*Proof*

From Lemma 1d we have *K*^⊥^(*B, z*^{*π*(*C*)}^) = *PK*^⊥^(*A, z*^{*C*}^). Letting *X* = *K*^⊥^(*A, z*^{*C*}^), we have:

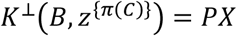

Substituting into the definition of *Q*_*X*_, the row-permutation applied by *s*, we have:

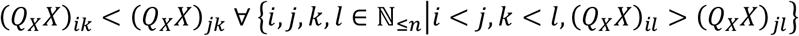

Multiplying internally by *P*^*T*^*P*, we have:

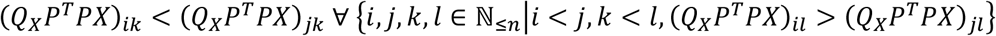

Factorising by (*Q*_*X*_*P*^*T*^), we have:

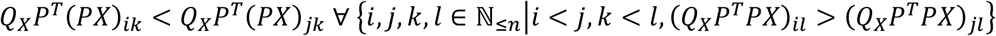

Thus, it is observed that *Q*_*X*_*P*^*T*^ is a solution to *Q*_*PX*_. Letting *Q*_*PX*_ = *Q*_*X*_*P*^*T*^, and multiplying from the right by *PX*, we have:

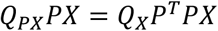

Thus:

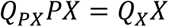

Substituting for the definitions of *s*(*PX*) and *s*(*X*), we have:

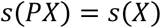

*End of Lemma 2*

**Lemma 3a** demonstrates that vertices with equal signatures are analogous. Thus, equality of ordered vertex signatures is sufficient to infer their analogy.

#### Lemma 3a.

If 𝒢_*A*_ ≅ 𝒢_*B*_ and *s* (*K*^⊥^(*A, z*^{*i*}^)) = *s* (*K*^⊥^(*B, z*^{*j*}^)) then 𝒢_*A*_(*𝓋*_*i*_) ↔ 𝒢_*B*_(*𝓋*_*j*_).

*Proof*

Let 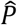 such that:

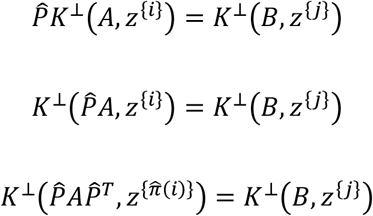

Expanding the Krylov matrices, we have:

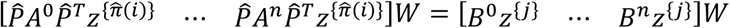

Thus 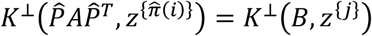. Deleting the left columns and factorising we have:

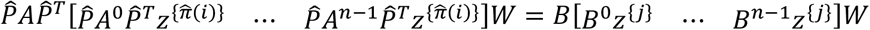

As 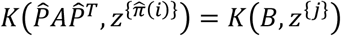 we also have:

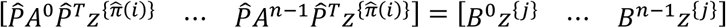

Deleting the right Krylov matrices, we 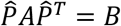. Therefore, there exists some 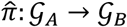 such that 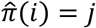. Hence, 𝒢_*A*_(*𝓋*_*i*_) ↔ 𝒢_*B*_(*𝓋*_*j*_).

*End of Lemma 3a*

**Lemma 3b** demonstrates that vertex sets with equal signatures are set analogous, extending the result of lemma 3a. Hence, equality of ordered vertex set signatures is sufficient to infer their set analogy.

#### Lemma 3b.

If 𝒢_*A*_ ≅ 𝒢_*B*_ and *s* (*K*(*A, z*^{*C*}^)) = *s* (*K*(*B, z*^{*D*}^)) then 𝒢_*A*_(*𝓋*_*C*_) ↔ 𝒢_*B*_(*𝓋*_*D*_).

*Proof*

As in lemma 3b, we have

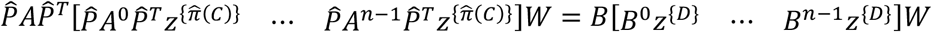

Therefore, there exists some 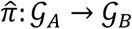 such that 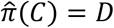

*End of Lemma 3b*

### Algorithm Overview

Combining the results of Lemma 2 and 3a it is observed that equality of ordered vertex signatures *s* (*K*^⊥^(*A, z*^{*i*}^)) and *s* (*K*^⊥^(*B, z*^{*j*}^)) is both necessary and sufficient to demonstrate analogy of vertices 𝒢_*A*_(*𝓋*_*i*_) and 𝒢_*B*_(*𝓋*_*j*_). Thus, the problem of vertex analogy testing is exactly linearised, with ℴ(*n*^3^) time complexity. For the first vertex in the source graph, 𝒢_*A*_(*𝓋*_1_), there exist *n* candidate vertices, 𝒢_*B*_(*𝓋*_*j*_), *j* ∈ {1, …, *n*}, which may be checked sequentially for analogy. Following identification of the first analogy 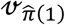, Lemma 3b may be employed to find analogous vertex sets. Vertex set analogies are constructed one vertex analogy at a time, so that all but one of the analogies in the set have already been established. Thus, the new analogy is evaluated whilst conditioning on the previously established analogies. For example, analogy to the vertex 2-set 𝒢_*A*_(*𝓋*_{1,2}_) is sought amongst the *n* − 1 candidate vertex 2-sets:

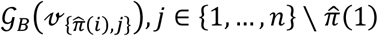

Likewise, analogy to the vertex set 𝒢_*A*_(*𝓋*_{1,…,|*C*|+1}_) is sought amongst the (*n* − |*C*|) candidate vertex sets:

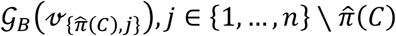

For the *n*th source vertex, all but one target vertex will have been assigned, and the final analogy may be deduced by exclusion. Therefore, within *n* − 1 iterations, an analogy will have been identified for each vertex in the source graph, yielding an isomorphic mapping if the assumption of isomorphism is true. Therefore, the algorithm is guaranteed to recover an isomorphic mapping if one exists. Figure 1 illustration the process of vertex signature computation and comparison in a toy application to a 3-vertex chain graph.

**Figure 1.**
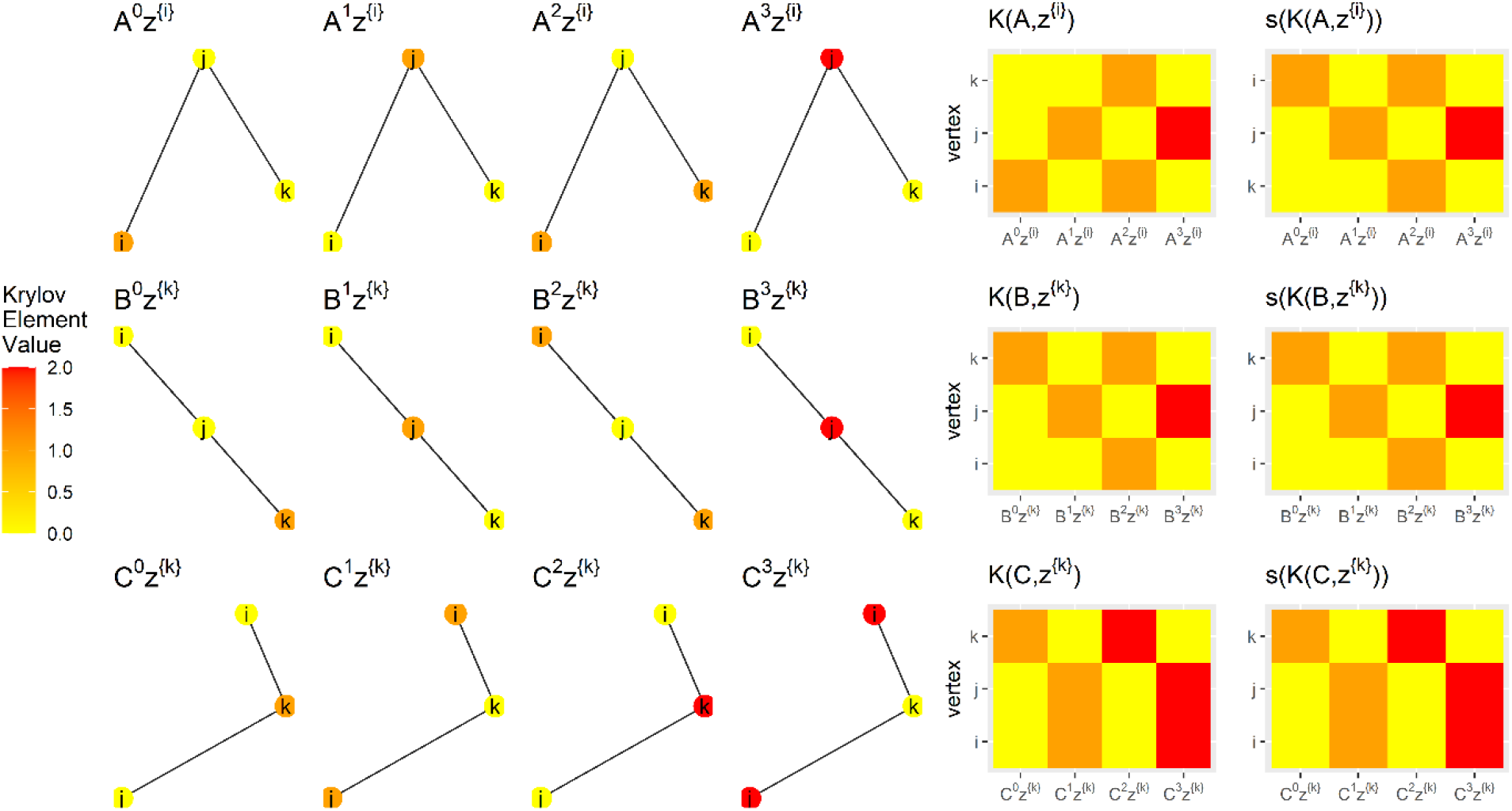
Illustration of the vertex analogy testing procedure in three isomorphic graphs, 𝒢_*A*_, 𝒢_*B*_ and 𝒢_*C*_. The Krylov matrix of the *i*th vertex in 𝒢_*A*_, *K*(*A, z*^{*i*}^) is generated by multiplying a binary indicator of the vertex position, *z*^{*i*}^, with adjacency matrix exponents. The matrix is row-ordered to give the ordered vertex signature, *s* (*K*(*A, z*^{*i*}^)). Equality of ordered vertex signatures indicates analogy of the corresponding vertices. Here 𝒢_*A*_(*𝓋*_*i*_) and 𝒢_*B*_(*𝓋*_*k*_) are analogous. In contrast, 𝒢_*A*_(*𝓋*_*i*_) and 𝒢_*C*_(*𝓋*_*k*_) are non-analogous, as their ordered vertex signatures differ.

#### Algorithm Pseudocode

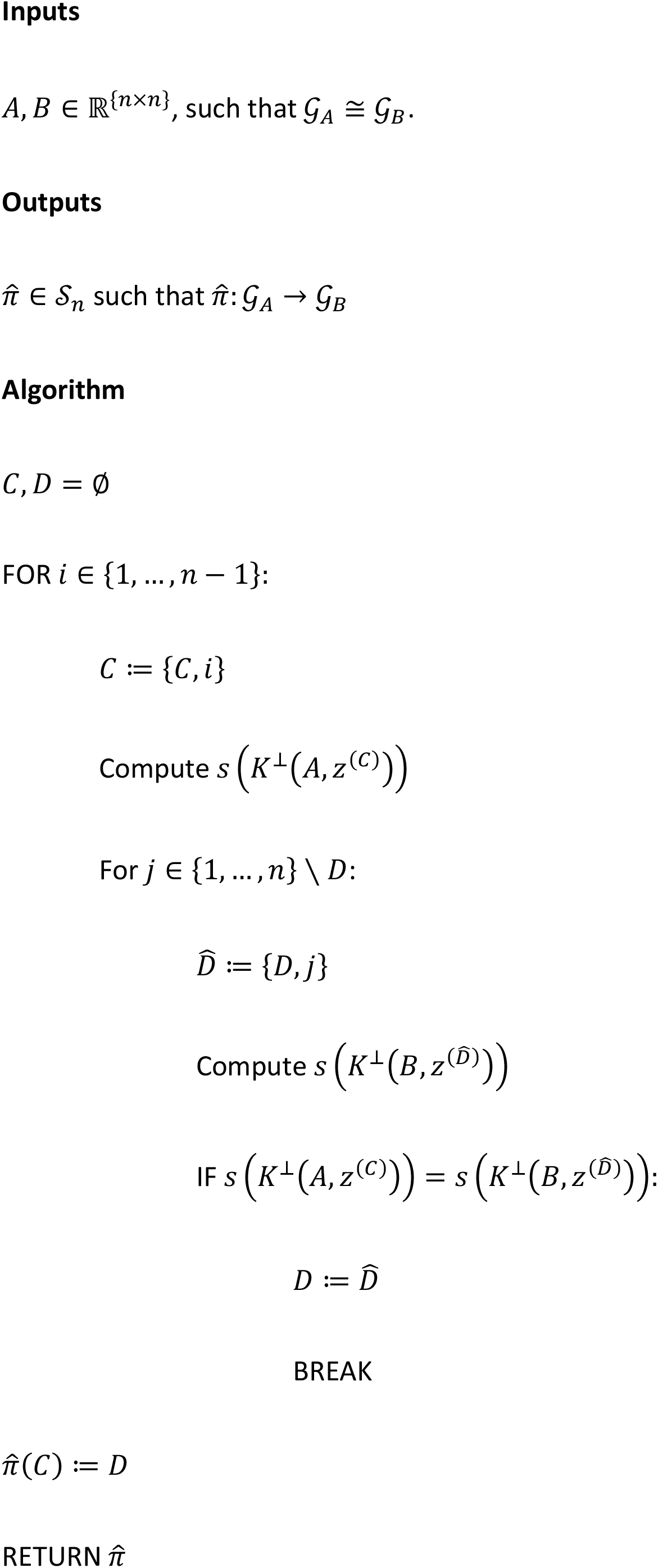

### Experiments

250 directed graphs were generated from a public repository of enzyme structures^13^. In each experiment, graph vertices were randomly permuted, and the algorithm applied to infer an isomorphic mapping. To alleviate numerical instability, Krylov matrix elements with absolute value < 10^−8^ were rounded to zero and column vectors were divided as appropriate to limit the absolute value of the maximum element at 10^8^. Where columns were orthogonalized to zero vectors, indicating complete subspace spanning^14^, no further columns were added to the Krylov matrices. For comparison, the BLISS heuristic graph matching algorithm^15^ was also applied. Code was written in the R language, with RStudio^16,17^. iGraph and abind libraries were employed^18,19^.

## Results

### Experimental Performance

Vertex analogy testing by our algorithm identified correct isomorphic mappings in every experiment (250/250, 100%). BLISS identified a incorrect isomorphic mappings in all but one experiment (1/250, 0.4%). All experimental results are provided in **Supplementary Table 1**.

### Complexity Analysis

Computation of orthogonalized Krylov matrices runs in ℴ(*n*^3^) time. For the first vertex in the source graph, *n* candidate vertex analogies exist in the target graph. By a process of deduction, if the graphs are isomorphic and the first *n* − 1 candidates are non-analogous, the *n*th vertex must be analogous. Hence, up to *n* − 1 orthogonal Krylov matrices must be computed and ordered, with the process stopping when an analogy is detected. For the second vertex in the source graph, only *n* − 1 target candidates remain, requiring computation of *n* − 2 ordered orthogonal Krylov matrices. Thus, in the worst case, all vertex analogies will be detected after computation of 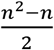 ordered vertex set signatures. Therefore, an isomorphic mapping may be identified in ℴ(*n*^5^) time in the worst case.

## Discussion

Theoretical analysis and experimental applications demonstrate that the algorithm recovers isomorphic mappings between small, directed graphs in polynomial-time. If exact arithmetic was available, this result would extend to large graphs. However, the algorithm’s dependence on recursive matrix exponentiation and orthogonalization exposes vulnerabilities to related to numerical precision. In mild cases, this limitation may be partially alleviated with appropriate rounding and numerical tolerance. Recursive exponentiation may also cause either vanishing or explosive growth of Krylov matrix elements. This problem may be addressed in some cases by normalising vectors.

The algorithm’s numerical instability and computational complexity limit applicability to large graphs. Accordingly, the proof-of-concept experiments presented here are limited to small graphs. Approaches for stabilising the algorithm in large graphs remain a topic for future research.

In contrast to previously presented spectral approaches ^3,8^, this algorithm does not involve graph perturbation, relying instead on distinctness of elements of the Krylov initialisation vector. The algorithm is guaranteed to recover an isomorphic mapping if one exists. If multiple mappings exist, a single solution will be returned, However, all mappings may be recovered systematically, by continuing to search for analogies beyond the first hit.

The algorithm provides a theoretical framework by which isomorphic mappings may be inferred between directed graphs, regardless of symmetry or degeneracy. Hence, an improvement in scope is offered on recent polynomial time approaches for undirected graphs^3^. As any real square matrix may be represented by a directed graph, this is the most generalisable approach to infer isomorphic mappings exactly. The theory underlying the algorithm yields insight into automorphism groups.

## Acknowledgements

The author would like to acknowledge the advice of Reimer Kuhn and Stefan Izaak at the Department of Theoretical Physics, King’s College London. The author would like to acknowledge the support of Vicky Goh and Gary Cook at the Department of Cancer Imaging, King’s College London; and Sophia Tsoka at the Department of Informatics, King’s College London.

## Availability of Code and Materials

All graph datasets included in this article were downloaded from a public repository^13^. All code required to reproduce this analysis is implemented in the R language and provided at https://github.com/robertoshea/graph_isomorphism_directed.

