## supplementary table 1 for "Exact polynomial-time isomorphism testing in directed graphs through comparison of vertex signatures in Krylov subspaces"

**Supplementary Table 1**. Experimental performance of each graph matching algorithm. In each experiment, the graph’s vertices were randomly permuted to produce a pair of isomorphic graphs. The graph inference algorithms were applied to infer an isomorphic mapping.

| **Vertices** | **Edges** | **Method** | **Correct mapping** | **Time**  (secs) | **Graph** |
| --- | --- | --- | --- | --- | --- |
| 37 | 336 | Vertex Analogy Testing | TRUE | 0.09 | enzyme_g1 |
| 37 | 336 | BLISS | FALSE | 0.01 | enzyme_g1 |
| 23 | 204 | Vertex Analogy Testing | TRUE | 0.04 | enzyme_g2 |
| 23 | 204 | BLISS | FALSE | 0 | enzyme_g2 |
| 25 | 184 | Vertex Analogy Testing | TRUE | 0.03 | enzyme_g3 |
| 25 | 184 | BLISS | FALSE | 0 | enzyme_g3 |
| 24 | 180 | Vertex Analogy Testing | TRUE | 0.02 | enzyme_g4 |
| 24 | 180 | BLISS | FALSE | 0 | enzyme_g4 |
| 23 | 180 | Vertex Analogy Testing | TRUE | 0.03 | enzyme_g5 |
| 23 | 180 | BLISS | FALSE | 0 | enzyme_g5 |
| 24 | 184 | Vertex Analogy Testing | TRUE | 0.03 | enzyme_g6 |
| 24 | 184 | BLISS | FALSE | 0 | enzyme_g6 |
| 26 | 236 | Vertex Analogy Testing | TRUE | 0.03 | enzyme_g7 |
| 26 | 236 | BLISS | FALSE | 0 | enzyme_g7 |
| 88 | 532 | Vertex Analogy Testing | TRUE | 0.26 | enzyme_g8 |
| 88 | 532 | BLISS | FALSE | 0 | enzyme_g8 |
| 23 | 156 | Vertex Analogy Testing | TRUE | 0.02 | enzyme_g9 |
| 23 | 156 | BLISS | FALSE | 0 | enzyme_g9 |
| 32 | 212 | Vertex Analogy Testing | TRUE | 0.04 | enzyme_g10 |
| 32 | 212 | BLISS | FALSE | 0 | enzyme_g10 |
| 4 | 24 | Vertex Analogy Testing | TRUE | 0 | enzyme_g11 |
| 4 | 24 | BLISS | FALSE | 0 | enzyme_g11 |
| 14 | 112 | Vertex Analogy Testing | TRUE | 0.01 | enzyme_g12 |
| 14 | 112 | BLISS | FALSE | 0 | enzyme_g12 |
| 42 | 300 | Vertex Analogy Testing | TRUE | 0.07 | enzyme_g13 |
| 42 | 300 | BLISS | FALSE | 0 | enzyme_g13 |
| 41 | 292 | Vertex Analogy Testing | TRUE | 0.04 | enzyme_g14 |
| 41 | 292 | BLISS | FALSE | 0 | enzyme_g14 |
| 36 | 256 | Vertex Analogy Testing | TRUE | 0.05 | enzyme_g15 |
| 36 | 256 | BLISS | FALSE | 0 | enzyme_g15 |
| 55 | 388 | Vertex Analogy Testing | TRUE | 0.12 | enzyme_g16 |
| 55 | 388 | BLISS | FALSE | 0 | enzyme_g16 |
| 40 | 380 | Vertex Analogy Testing | TRUE | 0.05 | enzyme_g17 |
| 40 | 380 | BLISS | FALSE | 0 | enzyme_g17 |
| 38 | 364 | Vertex Analogy Testing | TRUE | 0.05 | enzyme_g18 |
| 38 | 364 | BLISS | FALSE | 0 | enzyme_g18 |
| 2 | 4 | Vertex Analogy Testing | TRUE | 0 | enzyme_g19 |
| 2 | 4 | BLISS | TRUE | 0 | enzyme_g19 |
| 35 | 280 | Vertex Analogy Testing | TRUE | 0.04 | enzyme_g20 |
| 35 | 280 | BLISS | FALSE | 0 | enzyme_g20 |
| 42 | 360 | Vertex Analogy Testing | TRUE | 0.06 | enzyme_g21 |
| 42 | 360 | BLISS | FALSE | 0 | enzyme_g21 |
| 41 | 360 | Vertex Analogy Testing | TRUE | 0.06 | enzyme_g22 |
| 41 | 360 | BLISS | FALSE | 0 | enzyme_g22 |
| 39 | 324 | Vertex Analogy Testing | TRUE | 0.07 | enzyme_g23 |
| 39 | 324 | BLISS | FALSE | 0 | enzyme_g23 |
| 42 | 368 | Vertex Analogy Testing | TRUE | 0.06 | enzyme_g24 |
| 42 | 368 | BLISS | FALSE | 0 | enzyme_g24 |
| 41 | 352 | Vertex Analogy Testing | TRUE | 0.06 | enzyme_g25 |
| 41 | 352 | BLISS | FALSE | 0 | enzyme_g25 |
| 40 | 340 | Vertex Analogy Testing | TRUE | 0.06 | enzyme_g26 |
| 40 | 340 | BLISS | FALSE | 0 | enzyme_g26 |
| 37 | 332 | Vertex Analogy Testing | TRUE | 0.07 | enzyme_g27 |
| 37 | 332 | BLISS | FALSE | 0 | enzyme_g27 |
| 23 | 196 | Vertex Analogy Testing | TRUE | 0.03 | enzyme_g28 |
| 23 | 196 | BLISS | FALSE | 0 | enzyme_g28 |
| 22 | 188 | Vertex Analogy Testing | TRUE | 0.03 | enzyme_g29 |
| 22 | 188 | BLISS | FALSE | 0 | enzyme_g29 |
| 34 | 264 | Vertex Analogy Testing | TRUE | 0.05 | enzyme_g30 |
| 34 | 264 | BLISS | FALSE | 0 | enzyme_g30 |
| 38 | 324 | Vertex Analogy Testing | TRUE | 0.06 | enzyme_g31 |
| 38 | 324 | BLISS | FALSE | 0 | enzyme_g31 |
| 38 | 316 | Vertex Analogy Testing | TRUE | 0.06 | enzyme_g32 |
| 38 | 316 | BLISS | FALSE | 0 | enzyme_g32 |
| 39 | 328 | Vertex Analogy Testing | TRUE | 0.05 | enzyme_g33 |
| 39 | 328 | BLISS | FALSE | 0 | enzyme_g33 |
| 8 | 56 | Vertex Analogy Testing | TRUE | 0.01 | enzyme_g34 |
| 8 | 56 | BLISS | FALSE | 0 | enzyme_g34 |
| 23 | 176 | Vertex Analogy Testing | TRUE | 0.02 | enzyme_g35 |
| 23 | 176 | BLISS | FALSE | 0 | enzyme_g35 |
| 42 | 324 | Vertex Analogy Testing | TRUE | 0.07 | enzyme_g36 |
| 42 | 324 | BLISS | FALSE | 0 | enzyme_g36 |
| 42 | 320 | Vertex Analogy Testing | TRUE | 0.07 | enzyme_g37 |
| 42 | 320 | BLISS | FALSE | 0 | enzyme_g37 |
| 98 | 64 | Vertex Analogy Testing | TRUE | 0.2 | enzyme_g38 |
| 98 | 64 | BLISS | FALSE | 0 | enzyme_g38 |
| 24 | 172 | Vertex Analogy Testing | TRUE | 0.02 | enzyme_g39 |
| 24 | 172 | BLISS | FALSE | 0 | enzyme_g39 |
| 24 | 200 | Vertex Analogy Testing | TRUE | 0.03 | enzyme_g40 |
| 24 | 200 | BLISS | FALSE | 0 | enzyme_g40 |
| 47 | 380 | Vertex Analogy Testing | TRUE | 0.11 | enzyme_g41 |
| 47 | 380 | BLISS | FALSE | 0 | enzyme_g41 |
| 45 | 388 | Vertex Analogy Testing | TRUE | 0.08 | enzyme_g42 |
| 45 | 388 | BLISS | FALSE | 0.01 | enzyme_g42 |
| 45 | 380 | Vertex Analogy Testing | TRUE | 0.07 | enzyme_g43 |
| 45 | 380 | BLISS | FALSE | 0 | enzyme_g43 |
| 45 | 380 | Vertex Analogy Testing | TRUE | 0.06 | enzyme_g44 |
| 45 | 380 | BLISS | FALSE | 0 | enzyme_g44 |
| 46 | 388 | Vertex Analogy Testing | TRUE | 0.06 | enzyme_g45 |
| 46 | 388 | BLISS | FALSE | 0 | enzyme_g45 |
| 44 | 380 | Vertex Analogy Testing | TRUE | 0.08 | enzyme_g46 |
| 44 | 380 | BLISS | FALSE | 0 | enzyme_g46 |
| 30 | 204 | Vertex Analogy Testing | TRUE | 0.03 | enzyme_g47 |
| 30 | 204 | BLISS | FALSE | 0 | enzyme_g47 |
| 32 | 228 | Vertex Analogy Testing | TRUE | 0.04 | enzyme_g48 |
| 32 | 228 | BLISS | FALSE | 0 | enzyme_g48 |
| 33 | 256 | Vertex Analogy Testing | TRUE | 0.04 | enzyme_g49 |
| 33 | 256 | BLISS | FALSE | 0 | enzyme_g49 |
| 9 | 60 | Vertex Analogy Testing | TRUE | 0.01 | enzyme_g50 |
| 9 | 60 | BLISS | FALSE | 0 | enzyme_g50 |
| 27 | 188 | Vertex Analogy Testing | TRUE | 0.03 | enzyme_g51 |
| 27 | 188 | BLISS | FALSE | 0 | enzyme_g51 |
| 39 | 340 | Vertex Analogy Testing | TRUE | 0.08 | enzyme_g52 |
| 39 | 340 | BLISS | FALSE | 0 | enzyme_g52 |
| 16 | 124 | Vertex Analogy Testing | TRUE | 0.02 | enzyme_g53 |
| 16 | 124 | BLISS | FALSE | 0 | enzyme_g53 |
| 18 | 124 | Vertex Analogy Testing | TRUE | 0.02 | enzyme_g54 |
| 18 | 124 | BLISS | FALSE | 0 | enzyme_g54 |
| 7 | 48 | Vertex Analogy Testing | TRUE | 0 | enzyme_g55 |
| 7 | 48 | BLISS | FALSE | 0 | enzyme_g55 |
| 18 | 148 | Vertex Analogy Testing | TRUE | 0.03 | enzyme_g56 |
| 18 | 148 | BLISS | FALSE | 0 | enzyme_g56 |
| 10 | 72 | Vertex Analogy Testing | TRUE | 0.01 | enzyme_g57 |
| 10 | 72 | BLISS | FALSE | 0 | enzyme_g57 |
| 21 | 200 | Vertex Analogy Testing | TRUE | 0.02 | enzyme_g58 |
| 21 | 200 | BLISS | FALSE | 0 | enzyme_g58 |
| 18 | 128 | Vertex Analogy Testing | TRUE | 0.02 | enzyme_g59 |
| 18 | 128 | BLISS | FALSE | 0 | enzyme_g59 |
| 10 | 72 | Vertex Analogy Testing | TRUE | 0.01 | enzyme_g60 |
| 10 | 72 | BLISS | FALSE | 0 | enzyme_g60 |
| 9 | 64 | Vertex Analogy Testing | TRUE | 0.01 | enzyme_g61 |
| 9 | 64 | BLISS | FALSE | 0 | enzyme_g61 |
| 39 | 280 | Vertex Analogy Testing | TRUE | 0.07 | enzyme_g62 |
| 39 | 280 | BLISS | FALSE | 0 | enzyme_g62 |
| 33 | 220 | Vertex Analogy Testing | TRUE | 0.05 | enzyme_g63 |
| 33 | 220 | BLISS | FALSE | 0 | enzyme_g63 |
| 29 | 196 | Vertex Analogy Testing | TRUE | 0.03 | enzyme_g64 |
| 29 | 196 | BLISS | FALSE | 0 | enzyme_g64 |
| 24 | 156 | Vertex Analogy Testing | TRUE | 0.03 | enzyme_g65 |
| 24 | 156 | BLISS | FALSE | 0 | enzyme_g65 |
| 25 | 172 | Vertex Analogy Testing | TRUE | 0.03 | enzyme_g66 |
| 25 | 172 | BLISS | FALSE | 0 | enzyme_g66 |
| 30 | 216 | Vertex Analogy Testing | TRUE | 0.04 | enzyme_g67 |
| 30 | 216 | BLISS | FALSE | 0 | enzyme_g67 |
| 38 | 300 | Vertex Analogy Testing | TRUE | 0.06 | enzyme_g68 |
| 38 | 300 | BLISS | FALSE | 0 | enzyme_g68 |
| 28 | 220 | Vertex Analogy Testing | TRUE | 0.04 | enzyme_g69 |
| 28 | 220 | BLISS | FALSE | 0 | enzyme_g69 |
| 28 | 216 | Vertex Analogy Testing | TRUE | 0.04 | enzyme_g70 |
| 28 | 216 | BLISS | FALSE | 0 | enzyme_g70 |
| 38 | 352 | Vertex Analogy Testing | TRUE | 0.06 | enzyme_g71 |
| 38 | 352 | BLISS | FALSE | 0 | enzyme_g71 |
| 40 | 280 | Vertex Analogy Testing | TRUE | 0.09 | enzyme_g72 |
| 40 | 280 | BLISS | FALSE | 0 | enzyme_g72 |
| 40 | 348 | Vertex Analogy Testing | TRUE | 0.06 | enzyme_g73 |
| 40 | 348 | BLISS | FALSE | 0 | enzyme_g73 |
| 42 | 376 | Vertex Analogy Testing | TRUE | 0.07 | enzyme_g74 |
| 42 | 376 | BLISS | FALSE | 0 | enzyme_g74 |
| 20 | 144 | Vertex Analogy Testing | TRUE | 0.03 | enzyme_g75 |
| 20 | 144 | BLISS | FALSE | 0 | enzyme_g75 |
| 19 | 140 | Vertex Analogy Testing | TRUE | 0.03 | enzyme_g76 |
| 19 | 140 | BLISS | FALSE | 0 | enzyme_g76 |
| 16 | 112 | Vertex Analogy Testing | TRUE | 0.03 | enzyme_g77 |
| 16 | 112 | BLISS | FALSE | 0 | enzyme_g77 |
| 17 | 116 | Vertex Analogy Testing | TRUE | 0.02 | enzyme_g78 |
| 17 | 116 | BLISS | FALSE | 0 | enzyme_g78 |
| 20 | 160 | Vertex Analogy Testing | TRUE | 0.03 | enzyme_g79 |
| 20 | 160 | BLISS | FALSE | 0 | enzyme_g79 |
| 18 | 128 | Vertex Analogy Testing | TRUE | 0.03 | enzyme_g80 |
| 18 | 128 | BLISS | FALSE | 0 | enzyme_g80 |
| 33 | 276 | Vertex Analogy Testing | TRUE | 0.04 | enzyme_g81 |
| 33 | 276 | BLISS | FALSE | 0 | enzyme_g81 |
| 23 | 176 | Vertex Analogy Testing | TRUE | 0.04 | enzyme_g82 |
| 23 | 176 | BLISS | FALSE | 0 | enzyme_g82 |
| 23 | 168 | Vertex Analogy Testing | TRUE | 0.03 | enzyme_g83 |
| 23 | 168 | BLISS | FALSE | 0 | enzyme_g83 |
| 35 | 296 | Vertex Analogy Testing | TRUE | 0.05 | enzyme_g84 |
| 35 | 296 | BLISS | FALSE | 0 | enzyme_g84 |
| 33 | 272 | Vertex Analogy Testing | TRUE | 0.05 | enzyme_g85 |
| 33 | 272 | BLISS | FALSE | 0 | enzyme_g85 |
| 39 | 276 | Vertex Analogy Testing | TRUE | 0.05 | enzyme_g86 |
| 39 | 276 | BLISS | FALSE | 0 | enzyme_g86 |
| 40 | 292 | Vertex Analogy Testing | TRUE | 0.06 | enzyme_g87 |
| 40 | 292 | BLISS | FALSE | 0 | enzyme_g87 |
| 38 | 364 | Vertex Analogy Testing | TRUE | 0.07 | enzyme_g88 |
| 38 | 364 | BLISS | FALSE | 0 | enzyme_g88 |
| 40 | 376 | Vertex Analogy Testing | TRUE | 0.06 | enzyme_g89 |
| 40 | 376 | BLISS | FALSE | 0 | enzyme_g89 |
| 38 | 264 | Vertex Analogy Testing | TRUE | 0.05 | enzyme_g90 |
| 38 | 264 | BLISS | FALSE | 0 | enzyme_g90 |
| 37 | 316 | Vertex Analogy Testing | TRUE | 0.06 | enzyme_g91 |
| 37 | 316 | BLISS | FALSE | 0 | enzyme_g91 |
| 36 | 308 | Vertex Analogy Testing | TRUE | 0.07 | enzyme_g92 |
| 36 | 308 | BLISS | FALSE | 0 | enzyme_g92 |
| 34 | 224 | Vertex Analogy Testing | TRUE | 0.04 | enzyme_g93 |
| 34 | 224 | BLISS | FALSE | 0 | enzyme_g93 |
| 34 | 268 | Vertex Analogy Testing | TRUE | 0.04 | enzyme_g94 |
| 34 | 268 | BLISS | FALSE | 0 | enzyme_g94 |
| 32 | 256 | Vertex Analogy Testing | TRUE | 0.03 | enzyme_g95 |
| 32 | 256 | BLISS | FALSE | 0 | enzyme_g95 |
| 18 | 128 | Vertex Analogy Testing | TRUE | 0.02 | enzyme_g96 |
| 18 | 128 | BLISS | FALSE | 0 | enzyme_g96 |
| 32 | 260 | Vertex Analogy Testing | TRUE | 0.04 | enzyme_g97 |
| 32 | 260 | BLISS | FALSE | 0 | enzyme_g97 |
| 34 | 268 | Vertex Analogy Testing | TRUE | 0.05 | enzyme_g98 |
| 34 | 268 | BLISS | FALSE | 0 | enzyme_g98 |
| 30 | 232 | Vertex Analogy Testing | TRUE | 0.04 | enzyme_g99 |
| 30 | 232 | BLISS | FALSE | 0 | enzyme_g99 |
| 5 | 36 | Vertex Analogy Testing | TRUE | 0 | enzyme_g100 |
| 5 | 36 | BLISS | FALSE | 0 | enzyme_g100 |
| 45 | 352 | Vertex Analogy Testing | TRUE | 0.06 | enzyme_g101 |
| 45 | 352 | BLISS | FALSE | 0 | enzyme_g101 |
| 42 | 328 | Vertex Analogy Testing | TRUE | 0.05 | enzyme_g102 |
| 42 | 328 | BLISS | FALSE | 0 | enzyme_g102 |
| 59 | 460 | Vertex Analogy Testing | TRUE | 0.1 | enzyme_g103 |
| 59 | 460 | BLISS | FALSE | 0 | enzyme_g103 |
| 32 | 256 | Vertex Analogy Testing | TRUE | 0.03 | enzyme_g104 |
| 32 | 256 | BLISS | FALSE | 0 | enzyme_g104 |
| 33 | 276 | Vertex Analogy Testing | TRUE | 0.04 | enzyme_g105 |
| 33 | 276 | BLISS | FALSE | 0 | enzyme_g105 |
| 17 | 120 | Vertex Analogy Testing | TRUE | 0.03 | enzyme_g106 |
| 17 | 120 | BLISS | FALSE | 0 | enzyme_g106 |
| 39 | 332 | Vertex Analogy Testing | TRUE | 0.06 | enzyme_g107 |
| 39 | 332 | BLISS | FALSE | 0 | enzyme_g107 |
| 38 | 328 | Vertex Analogy Testing | TRUE | 0.08 | enzyme_g108 |
| 38 | 328 | BLISS | FALSE | 0 | enzyme_g108 |
| 35 | 280 | Vertex Analogy Testing | TRUE | 0.05 | enzyme_g109 |
| 35 | 280 | BLISS | FALSE | 0 | enzyme_g109 |
| 16 | 116 | Vertex Analogy Testing | TRUE | 0.02 | enzyme_g110 |
| 16 | 116 | BLISS | FALSE | 0 | enzyme_g110 |
| 26 | 196 | Vertex Analogy Testing | TRUE | 0.04 | enzyme_g111 |
| 26 | 196 | BLISS | FALSE | 0 | enzyme_g111 |
| 51 | 380 | Vertex Analogy Testing | TRUE | 0.08 | enzyme_g112 |
| 51 | 380 | BLISS | FALSE | 0 | enzyme_g112 |
| 52 | 392 | Vertex Analogy Testing | TRUE | 0.1 | enzyme_g113 |
| 52 | 392 | BLISS | FALSE | 0 | enzyme_g113 |
| 25 | 188 | Vertex Analogy Testing | TRUE | 0.04 | enzyme_g114 |
| 25 | 188 | BLISS | FALSE | 0 | enzyme_g114 |
| 27 | 204 | Vertex Analogy Testing | TRUE | 0.03 | enzyme_g115 |
| 27 | 204 | BLISS | FALSE | 0 | enzyme_g115 |
| 42 | 296 | Vertex Analogy Testing | TRUE | 0.04 | enzyme_g116 |
| 42 | 296 | BLISS | FALSE | 0 | enzyme_g116 |
| 46 | 360 | Vertex Analogy Testing | TRUE | 0.09 | enzyme_g117 |
| 46 | 360 | BLISS | FALSE | 0 | enzyme_g117 |
| 96 | 484 | Vertex Analogy Testing | TRUE | 0.4 | enzyme_g118 |
| 96 | 484 | BLISS | FALSE | 0 | enzyme_g118 |
| 12 | 88 | Vertex Analogy Testing | TRUE | 0.01 | enzyme_g119 |
| 12 | 88 | BLISS | FALSE | 0 | enzyme_g119 |
| 22 | 184 | Vertex Analogy Testing | TRUE | 0.03 | enzyme_g120 |
| 22 | 184 | BLISS | FALSE | 0 | enzyme_g120 |
| 42 | 328 | Vertex Analogy Testing | TRUE | 0.07 | enzyme_g121 |
| 42 | 328 | BLISS | FALSE | 0 | enzyme_g121 |
| 14 | 116 | Vertex Analogy Testing | TRUE | 0.01 | enzyme_g122 |
| 14 | 116 | BLISS | FALSE | 0 | enzyme_g122 |
| 90 | 508 | Vertex Analogy Testing | TRUE | 0.29 | enzyme_g123 |
| 90 | 508 | BLISS | FALSE | 0 | enzyme_g123 |
| 14 | 124 | Vertex Analogy Testing | TRUE | 0.01 | enzyme_g124 |
| 14 | 124 | BLISS | FALSE | 0 | enzyme_g124 |
| 14 | 132 | Vertex Analogy Testing | TRUE | 0.02 | enzyme_g125 |
| 14 | 132 | BLISS | FALSE | 0 | enzyme_g125 |
| 32 | 188 | Vertex Analogy Testing | TRUE | 0.06 | enzyme_g126 |
| 32 | 188 | BLISS | FALSE | 0 | enzyme_g126 |
| 11 | 92 | Vertex Analogy Testing | TRUE | 0.01 | enzyme_g127 |
| 11 | 92 | BLISS | FALSE | 0 | enzyme_g127 |
| 26 | 148 | Vertex Analogy Testing | TRUE | 0.04 | enzyme_g128 |
| 26 | 148 | BLISS | FALSE | 0 | enzyme_g128 |
| 11 | 80 | Vertex Analogy Testing | TRUE | 0.01 | enzyme_g129 |
| 11 | 80 | BLISS | FALSE | 0 | enzyme_g129 |
| 14 | 100 | Vertex Analogy Testing | TRUE | 0.01 | enzyme_g130 |
| 14 | 100 | BLISS | FALSE | 0 | enzyme_g130 |
| 18 | 140 | Vertex Analogy Testing | TRUE | 0.02 | enzyme_g131 |
| 18 | 140 | BLISS | FALSE | 0 | enzyme_g131 |
| 16 | 128 | Vertex Analogy Testing | TRUE | 0.02 | enzyme_g132 |
| 16 | 128 | BLISS | FALSE | 0 | enzyme_g132 |
| 17 | 136 | Vertex Analogy Testing | TRUE | 0.01 | enzyme_g133 |
| 17 | 136 | BLISS | FALSE | 0 | enzyme_g133 |
| 32 | 204 | Vertex Analogy Testing | TRUE | 0.06 | enzyme_g134 |
| 32 | 204 | BLISS | FALSE | 0 | enzyme_g134 |
| 13 | 104 | Vertex Analogy Testing | TRUE | 0.01 | enzyme_g135 |
| 13 | 104 | BLISS | FALSE | 0 | enzyme_g135 |
| 3 | 12 | Vertex Analogy Testing | TRUE | 0 | enzyme_g136 |
| 3 | 12 | BLISS | FALSE | 0 | enzyme_g136 |
| 22 | 160 | Vertex Analogy Testing | TRUE | 0.03 | enzyme_g137 |
| 22 | 160 | BLISS | FALSE | 0 | enzyme_g137 |
| 16 | 144 | Vertex Analogy Testing | TRUE | 0.02 | enzyme_g138 |
| 16 | 144 | BLISS | FALSE | 0 | enzyme_g138 |
| 38 | 328 | Vertex Analogy Testing | TRUE | 0.06 | enzyme_g139 |
| 38 | 328 | BLISS | FALSE | 0 | enzyme_g139 |
| 13 | 112 | Vertex Analogy Testing | TRUE | 0.02 | enzyme_g140 |
| 13 | 112 | BLISS | FALSE | 0 | enzyme_g140 |
| 12 | 96 | Vertex Analogy Testing | TRUE | 0.02 | enzyme_g141 |
| 12 | 96 | BLISS | FALSE | 0 | enzyme_g141 |
| 14 | 116 | Vertex Analogy Testing | TRUE | 0.01 | enzyme_g142 |
| 14 | 116 | BLISS | FALSE | 0 | enzyme_g142 |
| 39 | 244 | Vertex Analogy Testing | TRUE | 0.08 | enzyme_g143 |
| 39 | 244 | BLISS | FALSE | 0 | enzyme_g143 |
| 19 | 140 | Vertex Analogy Testing | TRUE | 0.03 | enzyme_g144 |
| 19 | 140 | BLISS | FALSE | 0 | enzyme_g144 |
| 20 | 156 | Vertex Analogy Testing | TRUE | 0.02 | enzyme_g145 |
| 20 | 156 | BLISS | FALSE | 0 | enzyme_g145 |
| 39 | 348 | Vertex Analogy Testing | TRUE | 0.07 | enzyme_g146 |
| 39 | 348 | BLISS | FALSE | 0 | enzyme_g146 |
| 40 | 336 | Vertex Analogy Testing | TRUE | 0.06 | enzyme_g147 |
| 40 | 336 | BLISS | FALSE | 0 | enzyme_g147 |
| 39 | 320 | Vertex Analogy Testing | TRUE | 0.05 | enzyme_g148 |
| 39 | 320 | BLISS | FALSE | 0 | enzyme_g148 |
| 39 | 328 | Vertex Analogy Testing | TRUE | 0.06 | enzyme_g149 |
| 39 | 328 | BLISS | FALSE | 0 | enzyme_g149 |
| 29 | 164 | Vertex Analogy Testing | TRUE | 0.05 | enzyme_g150 |
| 29 | 164 | BLISS | FALSE | 0 | enzyme_g150 |
| 22 | 160 | Vertex Analogy Testing | TRUE | 0.03 | enzyme_g151 |
| 22 | 160 | BLISS | FALSE | 0 | enzyme_g151 |
| 11 | 88 | Vertex Analogy Testing | TRUE | 0.01 | enzyme_g152 |
| 11 | 88 | BLISS | FALSE | 0 | enzyme_g152 |
| 8 | 64 | Vertex Analogy Testing | TRUE | 0.01 | enzyme_g153 |
| 8 | 64 | BLISS | FALSE | 0 | enzyme_g153 |
| 13 | 88 | Vertex Analogy Testing | TRUE | 0.01 | enzyme_g154 |
| 13 | 88 | BLISS | FALSE | 0 | enzyme_g154 |
| 18 | 132 | Vertex Analogy Testing | TRUE | 0.02 | enzyme_g155 |
| 18 | 132 | BLISS | FALSE | 0 | enzyme_g155 |
| 12 | 88 | Vertex Analogy Testing | TRUE | 0.01 | enzyme_g156 |
| 12 | 88 | BLISS | FALSE | 0 | enzyme_g156 |
| 8 | 60 | Vertex Analogy Testing | TRUE | 0.01 | enzyme_g157 |
| 8 | 60 | BLISS | FALSE | 0 | enzyme_g157 |
| 40 | 252 | Vertex Analogy Testing | TRUE | 0.07 | enzyme_g158 |
| 40 | 252 | BLISS | FALSE | 0 | enzyme_g158 |
| 12 | 100 | Vertex Analogy Testing | TRUE | 0.01 | enzyme_g159 |
| 12 | 100 | BLISS | FALSE | 0 | enzyme_g159 |
| 22 | 192 | Vertex Analogy Testing | TRUE | 0.02 | enzyme_g160 |
| 22 | 192 | BLISS | FALSE | 0 | enzyme_g160 |
| 22 | 168 | Vertex Analogy Testing | TRUE | 0.02 | enzyme_g161 |
| 22 | 168 | BLISS | FALSE | 0 | enzyme_g161 |
| 18 | 128 | Vertex Analogy Testing | TRUE | 0.02 | enzyme_g162 |
| 18 | 128 | BLISS | FALSE | 0 | enzyme_g162 |
| 12 | 88 | Vertex Analogy Testing | TRUE | 0.02 | enzyme_g163 |
| 12 | 88 | BLISS | FALSE | 0 | enzyme_g163 |
| 17 | 144 | Vertex Analogy Testing | TRUE | 0.02 | enzyme_g164 |
| 17 | 144 | BLISS | FALSE | 0 | enzyme_g164 |
| 14 | 104 | Vertex Analogy Testing | TRUE | 0.01 | enzyme_g165 |
| 14 | 104 | BLISS | FALSE | 0 | enzyme_g165 |
| 22 | 144 | Vertex Analogy Testing | TRUE | 0.02 | enzyme_g166 |
| 22 | 144 | BLISS | FALSE | 0 | enzyme_g166 |
| 43 | 376 | Vertex Analogy Testing | TRUE | 0.07 | enzyme_g167 |
| 43 | 376 | BLISS | FALSE | 0 | enzyme_g167 |
| 42 | 336 | Vertex Analogy Testing | TRUE | 0.05 | enzyme_g168 |
| 42 | 336 | BLISS | FALSE | 0 | enzyme_g168 |
| 44 | 376 | Vertex Analogy Testing | TRUE | 0.06 | enzyme_g169 |
| 44 | 376 | BLISS | FALSE | 0 | enzyme_g169 |
| 24 | 164 | Vertex Analogy Testing | TRUE | 0.03 | enzyme_g170 |
| 24 | 164 | BLISS | FALSE | 0 | enzyme_g170 |
| 48 | 396 | Vertex Analogy Testing | TRUE | 0.09 | enzyme_g171 |
| 48 | 396 | BLISS | FALSE | 0 | enzyme_g171 |
| 25 | 184 | Vertex Analogy Testing | TRUE | 0.03 | enzyme_g172 |
| 25 | 184 | BLISS | FALSE | 0 | enzyme_g172 |
| 46 | 384 | Vertex Analogy Testing | TRUE | 0.06 | enzyme_g173 |
| 46 | 384 | BLISS | FALSE | 0 | enzyme_g173 |
| 48 | 392 | Vertex Analogy Testing | TRUE | 0.11 | enzyme_g174 |
| 48 | 392 | BLISS | FALSE | 0 | enzyme_g174 |
| 25 | 172 | Vertex Analogy Testing | TRUE | 0.02 | enzyme_g175 |
| 25 | 172 | BLISS | FALSE | 0 | enzyme_g175 |
| 48 | 384 | Vertex Analogy Testing | TRUE | 0.07 | enzyme_g176 |
| 48 | 384 | BLISS | FALSE | 0 | enzyme_g176 |
| 44 | 384 | Vertex Analogy Testing | TRUE | 0.09 | enzyme_g177 |
| 44 | 384 | BLISS | FALSE | 0 | enzyme_g177 |
| 42 | 352 | Vertex Analogy Testing | TRUE | 0.07 | enzyme_g178 |
| 42 | 352 | BLISS | FALSE | 0 | enzyme_g178 |
| 40 | 236 | Vertex Analogy Testing | TRUE | 0.06 | enzyme_g179 |
| 40 | 236 | BLISS | FALSE | 0 | enzyme_g179 |
| 38 | 240 | Vertex Analogy Testing | TRUE | 0.06 | enzyme_g180 |
| 38 | 240 | BLISS | FALSE | 0 | enzyme_g180 |
| 40 | 324 | Vertex Analogy Testing | TRUE | 0.06 | enzyme_g181 |
| 40 | 324 | BLISS | FALSE | 0 | enzyme_g181 |
| 41 | 344 | Vertex Analogy Testing | TRUE | 0.06 | enzyme_g182 |
| 41 | 344 | BLISS | FALSE | 0 | enzyme_g182 |
| 42 | 376 | Vertex Analogy Testing | TRUE | 0.08 | enzyme_g183 |
| 42 | 376 | BLISS | FALSE | 0 | enzyme_g183 |
| 40 | 364 | Vertex Analogy Testing | TRUE | 0.08 | enzyme_g184 |
| 40 | 364 | BLISS | FALSE | 0 | enzyme_g184 |
| 44 | 380 | Vertex Analogy Testing | TRUE | 0.07 | enzyme_g185 |
| 44 | 380 | BLISS | FALSE | 0 | enzyme_g185 |
| 24 | 160 | Vertex Analogy Testing | TRUE | 0.03 | enzyme_g186 |
| 24 | 160 | BLISS | FALSE | 0 | enzyme_g186 |
| 44 | 376 | Vertex Analogy Testing | TRUE | 0.07 | enzyme_g187 |
| 44 | 376 | BLISS | FALSE | 0 | enzyme_g187 |
| 20 | 156 | Vertex Analogy Testing | TRUE | 0.02 | enzyme_g188 |
| 20 | 156 | BLISS | FALSE | 0 | enzyme_g188 |
| 42 | 380 | Vertex Analogy Testing | TRUE | 0.08 | enzyme_g189 |
| 42 | 380 | BLISS | FALSE | 0 | enzyme_g189 |
| 27 | 196 | Vertex Analogy Testing | TRUE | 0.04 | enzyme_g190 |
| 27 | 196 | BLISS | FALSE | 0 | enzyme_g190 |
| 48 | 384 | Vertex Analogy Testing | TRUE | 0.11 | enzyme_g191 |
| 48 | 384 | BLISS | FALSE | 0 | enzyme_g191 |
| 31 | 264 | Vertex Analogy Testing | TRUE | 0.06 | enzyme_g192 |
| 31 | 264 | BLISS | FALSE | 0 | enzyme_g192 |
| 30 | 240 | Vertex Analogy Testing | TRUE | 0.04 | enzyme_g193 |
| 30 | 240 | BLISS | FALSE | 0 | enzyme_g193 |
| 46 | 400 | Vertex Analogy Testing | TRUE | 0.09 | enzyme_g194 |
| 46 | 400 | BLISS | FALSE | 0 | enzyme_g194 |
| 47 | 368 | Vertex Analogy Testing | TRUE | 0.08 | enzyme_g195 |
| 47 | 368 | BLISS | FALSE | 0 | enzyme_g195 |
| 50 | 344 | Vertex Analogy Testing | TRUE | 0.08 | enzyme_g196 |
| 50 | 344 | BLISS | FALSE | 0 | enzyme_g196 |
| 40 | 308 | Vertex Analogy Testing | TRUE | 0.08 | enzyme_g197 |
| 40 | 308 | BLISS | FALSE | 0 | enzyme_g197 |
| 55 | 308 | Vertex Analogy Testing | TRUE | 0.09 | enzyme_g198 |
| 55 | 308 | BLISS | FALSE | 0 | enzyme_g198 |
| 62 | 432 | Vertex Analogy Testing | TRUE | 0.12 | enzyme_g199 |
| 62 | 432 | BLISS | FALSE | 0 | enzyme_g199 |
| 34 | 288 | Vertex Analogy Testing | TRUE | 0.05 | enzyme_g200 |
| 34 | 288 | BLISS | FALSE | 0 | enzyme_g200 |
| 29 | 212 | Vertex Analogy Testing | TRUE | 0.03 | enzyme_g201 |
| 29 | 212 | BLISS | FALSE | 0 | enzyme_g201 |
| 25 | 180 | Vertex Analogy Testing | TRUE | 0.03 | enzyme_g202 |
| 25 | 180 | BLISS | FALSE | 0 | enzyme_g202 |
| 56 | 400 | Vertex Analogy Testing | TRUE | 0.11 | enzyme_g203 |
| 56 | 400 | BLISS | FALSE | 0 | enzyme_g203 |
| 57 | 420 | Vertex Analogy Testing | TRUE | 0.14 | enzyme_g204 |
| 57 | 420 | BLISS | FALSE | 0 | enzyme_g204 |
| 27 | 244 | Vertex Analogy Testing | TRUE | 0.05 | enzyme_g205 |
| 27 | 244 | BLISS | FALSE | 0 | enzyme_g205 |
| 22 | 172 | Vertex Analogy Testing | TRUE | 0.03 | enzyme_g206 |
| 22 | 172 | BLISS | FALSE | 0 | enzyme_g206 |
| 24 | 172 | Vertex Analogy Testing | TRUE | 0.02 | enzyme_g207 |
| 24 | 172 | BLISS | FALSE | 0 | enzyme_g207 |
| 23 | 200 | Vertex Analogy Testing | TRUE | 0.04 | enzyme_g208 |
| 23 | 200 | BLISS | FALSE | 0 | enzyme_g208 |
| 57 | 404 | Vertex Analogy Testing | TRUE | 0.1 | enzyme_g209 |
| 57 | 404 | BLISS | FALSE | 0 | enzyme_g209 |
| 24 | 200 | Vertex Analogy Testing | TRUE | 0.03 | enzyme_g210 |
| 24 | 200 | BLISS | FALSE | 0 | enzyme_g210 |
| 24 | 200 | Vertex Analogy Testing | TRUE | 0.04 | enzyme_g211 |
| 24 | 200 | BLISS | FALSE | 0 | enzyme_g211 |
| 23 | 188 | Vertex Analogy Testing | TRUE | 0.03 | enzyme_g212 |
| 23 | 188 | BLISS | FALSE | 0 | enzyme_g212 |
| 25 | 220 | Vertex Analogy Testing | TRUE | 0.03 | enzyme_g213 |
| 25 | 220 | BLISS | FALSE | 0 | enzyme_g213 |
| 23 | 204 | Vertex Analogy Testing | TRUE | 0.04 | enzyme_g214 |
| 23 | 204 | BLISS | FALSE | 0 | enzyme_g214 |
| 48 | 416 | Vertex Analogy Testing | TRUE | 0.08 | enzyme_g215 |
| 48 | 416 | BLISS | FALSE | 0 | enzyme_g215 |
| 27 | 188 | Vertex Analogy Testing | TRUE | 0.04 | enzyme_g216 |
| 27 | 188 | BLISS | FALSE | 0 | enzyme_g216 |
| 15 | 136 | Vertex Analogy Testing | TRUE | 0.02 | enzyme_g217 |
| 15 | 136 | BLISS | FALSE | 0 | enzyme_g217 |
| 44 | 388 | Vertex Analogy Testing | TRUE | 0.09 | enzyme_g218 |
| 44 | 388 | BLISS | FALSE | 0 | enzyme_g218 |
| 29 | 256 | Vertex Analogy Testing | TRUE | 0.04 | enzyme_g219 |
| 29 | 256 | BLISS | FALSE | 0 | enzyme_g219 |
| 21 | 164 | Vertex Analogy Testing | TRUE | 0.02 | enzyme_g220 |
| 21 | 164 | BLISS | FALSE | 0 | enzyme_g220 |
| 34 | 252 | Vertex Analogy Testing | TRUE | 0.05 | enzyme_g221 |
| 34 | 252 | BLISS | FALSE | 0 | enzyme_g221 |
| 30 | 228 | Vertex Analogy Testing | TRUE | 0.04 | enzyme_g222 |
| 30 | 228 | BLISS | FALSE | 0 | enzyme_g222 |
| 40 | 316 | Vertex Analogy Testing | TRUE | 0.07 | enzyme_g223 |
| 40 | 316 | BLISS | FALSE | 0 | enzyme_g223 |
| 54 | 420 | Vertex Analogy Testing | TRUE | 0.11 | enzyme_g224 |
| 54 | 420 | BLISS | FALSE | 0 | enzyme_g224 |
| 18 | 140 | Vertex Analogy Testing | TRUE | 0.02 | enzyme_g225 |
| 18 | 140 | BLISS | FALSE | 0 | enzyme_g225 |
| 36 | 296 | Vertex Analogy Testing | TRUE | 0.05 | enzyme_g226 |
| 36 | 296 | BLISS | FALSE | 0 | enzyme_g226 |
| 37 | 324 | Vertex Analogy Testing | TRUE | 0.05 | enzyme_g227 |
| 37 | 324 | BLISS | FALSE | 0 | enzyme_g227 |
| 34 | 304 | Vertex Analogy Testing | TRUE | 0.05 | enzyme_g228 |
| 34 | 304 | BLISS | FALSE | 0 | enzyme_g228 |
| 23 | 200 | Vertex Analogy Testing | TRUE | 0.03 | enzyme_g229 |
| 23 | 200 | BLISS | FALSE | 0 | enzyme_g229 |
| 32 | 272 | Vertex Analogy Testing | TRUE | 0.04 | enzyme_g230 |
| 32 | 272 | BLISS | FALSE | 0 | enzyme_g230 |
| 33 | 304 | Vertex Analogy Testing | TRUE | 0.04 | enzyme_g231 |
| 33 | 304 | BLISS | FALSE | 0 | enzyme_g231 |
| 29 | 260 | Vertex Analogy Testing | TRUE | 0.03 | enzyme_g232 |
| 29 | 260 | BLISS | FALSE | 0 | enzyme_g232 |
| 39 | 284 | Vertex Analogy Testing | TRUE | 0.12 | enzyme_g233 |
| 39 | 284 | BLISS | FALSE | 0 | enzyme_g233 |
| 34 | 280 | Vertex Analogy Testing | TRUE | 0.04 | enzyme_g234 |
| 34 | 280 | BLISS | FALSE | 0 | enzyme_g234 |
| 36 | 284 | Vertex Analogy Testing | TRUE | 0.06 | enzyme_g235 |
| 36 | 284 | BLISS | FALSE | 0 | enzyme_g235 |
| 21 | 140 | Vertex Analogy Testing | TRUE | 0.03 | enzyme_g236 |
| 21 | 140 | BLISS | FALSE | 0 | enzyme_g236 |
| 6 | 44 | Vertex Analogy Testing | TRUE | 0 | enzyme_g237 |
| 6 | 44 | BLISS | FALSE | 0 | enzyme_g237 |
| 22 | 200 | Vertex Analogy Testing | TRUE | 0.03 | enzyme_g238 |
| 22 | 200 | BLISS | FALSE | 0 | enzyme_g238 |
| 18 | 160 | Vertex Analogy Testing | TRUE | 0.03 | enzyme_g239 |
| 18 | 160 | BLISS | FALSE | 0 | enzyme_g239 |
| 18 | 164 | Vertex Analogy Testing | TRUE | 0.02 | enzyme_g240 |
| 18 | 164 | BLISS | FALSE | 0 | enzyme_g240 |
| 34 | 256 | Vertex Analogy Testing | TRUE | 0.04 | enzyme_g241 |
| 34 | 256 | BLISS | FALSE | 0 | enzyme_g241 |
| 32 | 244 | Vertex Analogy Testing | TRUE | 0.03 | enzyme_g242 |
| 32 | 244 | BLISS | FALSE | 0 | enzyme_g242 |
| 33 | 248 | Vertex Analogy Testing | TRUE | 0.04 | enzyme_g243 |
| 33 | 248 | BLISS | FALSE | 0 | enzyme_g243 |
| 28 | 248 | Vertex Analogy Testing | TRUE | 0.03 | enzyme_g244 |
| 28 | 248 | BLISS | FALSE | 0 | enzyme_g244 |
| 40 | 352 | Vertex Analogy Testing | TRUE | 0.06 | enzyme_g245 |
| 40 | 352 | BLISS | FALSE | 0 | enzyme_g245 |
| 42 | 304 | Vertex Analogy Testing | TRUE | 0.05 | enzyme_g246 |
| 42 | 304 | BLISS | FALSE | 0 | enzyme_g246 |
| 39 | 232 | Vertex Analogy Testing | TRUE | 0.06 | enzyme_g247 |
| 39 | 232 | BLISS | FALSE | 0 | enzyme_g247 |
| 28 | 192 | Vertex Analogy Testing | TRUE | 0.03 | enzyme_g248 |
| 28 | 192 | BLISS | FALSE | 0 | enzyme_g248 |
| 29 | 208 | Vertex Analogy Testing | TRUE | 0.03 | enzyme_g249 |
| 29 | 208 | BLISS | FALSE | 0 | enzyme_g249 |
| 30 | 268 | Vertex Analogy Testing | TRUE | 0.05 | enzyme_g250 |
| 30 | 268 | BLISS | FALSE | 0 | enzyme_g250 |
